# Rapid precision targeting of nanoparticles to lung via caveolae pumping system in endothelium

**DOI:** 10.1101/2024.09.01.610705

**Authors:** Tapas R. Nayak, Adrian Chrastina, Jose Valencia, Robert Yedidsion, Tim Buss, Brittany Cederstrom, Jim Koziol, Michael D. Levin, Bogdan Olenyuk, Jan E. Schnitzer

**Author notes:** Corresponding Author: Dr. Jan Schnitzer Proteogenomics Research Institute for Systems Medicine 505 Coast Boulevard South La Jolla, CA 92037.

## Abstract

Modern medicine seeks precision targeting, imaging and therapy to maximize efficacy and avoid toxicities. Nanoparticles (NPs) have tremendous, yet unmet clinical potential to carry and deliver imaging and therapeutic agents systemically with tissue precision. But their size contributes to unwanted rapid scavenging by the reticulo-endothelial system (RES) and poor penetration of key endothelial cell (EC) barriers, both limiting target-tissue uptake, safety and efficacy. Here, we discover the extraordinary yet size-dependent ability of the EC caveolae pumping system (CPS) to deliver NPs rapidly and specifically into lungs. Gold and dendritic NPs are conjugated to aminopeptidase-P2 antibodies targeting caveolae of lung microvascular endothelium. SPECT-CT imaging and biodistribution analyses reveal that rat lungs extract most of the intravenously injected dose within minutes to achieve rapid blood clearance, high lung tissue concentrations well beyond peak blood levels, and precision lung imaging and targeting. Active transcytosis by caveolae greatly outperforms passive transvascular delivery and can even outpace RES scavenging. These results reveal how much ECs can both limit and promote tissue penetration of NPs and the power and limitations of the CPS. This study provides a new retargeting paradigm for small NPs to avoid RES uptake and achieve unprecedented rapid precision nanodelivery for future diagnostic and therapeutic applications.

---

The vast majority of deadly diseases occur in solid tissues that limit drug entry from the blood circulation and thus therapeutic efficacy. Systemic drug delivery and molecular medicine in general can benefit greatly from the capacity of NPs to carry and shield substantial payloads of various drugs and imaging probes ^1, 2, 3, 4, 5, 6^. NPs concentrate the drug in a protected environment that can improve the drug’s solubility, stability, pharmacokinetic (PK) and toxicity profiles^2, 3, 5, 7^. NPs can also be functionalized to seek and bind biomarkers specific to a desired cell type or tissue ^1, 3, 6, 8^. Yet their targeting precision and overall therapeutic benefit remains elusive because NP size: i) restricts entry into key tissues from the blood, leading to insufficient target tissue concentrations and ii) promotes undesired RES uptake, leading to lower blood levels, less drug for target tissues, and more toxicity, especially in RES tissues ^4, 6, 8^.

To reach its target molecule or cell inside a solid tissue for maximum efficacy, each NP must move from the blood microcirculation to the interstitium of the target-tissue by traversing the EC monolayer lining the microvasculature ^1, 6, 9^. All NPs, including those functionalized with very capable targeting ligands such as antibodies, rely on passive transvascular exchange driven mostly by diffusion and convection through open fenestrations, transient transendothelial channels, intercellular junctions and/or gaps between adjacent ECs ^6, 9^. Tissues with the leakiest blood vessels such as liver and other RES organs have a discontinuous endothelium with gaps readily permitting rapid NP entry and majority accumulation. But for all other tissues thwarted by more restrictive EC barriers and unhelpful diversion by RES scavenging, only minute fractions of the injected NP dose actually accumulate inside the target-tissue ^4, 6, 10^. The RES can clear the blood of NPs within minutes leaving few NPs to ever reach their intended target tissue through the slow and inefficient process of passive transvascular delivery. Methods to reduce or delay RES uptake have helped prolong circulation times, but precision-targeted tissue penetration remains the unmet challenge ^3, 4, 6, 10, 11, 12, 13, 14^.

After four decades of research and hundreds of studies, fundamental understanding of how NPs cross EC barriers remains poor ^6, 15^ and a median delivery of <2% of the NP dose actually ever reaches non-RES target-tissues including solid tumors ^10, 15, 16, 17^. Fueled by sparse clinical success, some scientists have even debated whether systemic nanodelivery is a futile therapeutic strategy^11, 12, 13, 14^. On the contrary, NPs readily cross EC barriers to enter RES tissues which can easily and rapidly accumulate the majority of the injected NP dose. Finally in the last decade, NP delivery companies that have survived years of challenging cancer clinical trials have switched to utilizing NP’s natural tissue tropism to develop systemic liver therapies with notable recent success ^18, 19^. New delivery strategies are urgently needed to allow the great promise of nanocarriers to be realized inside non-RES tissues.

Most tissues have blood vessels lined with continuous ECs that restrict NP penetration but have an abundant population of specialized plasmalemmal invaginations called caveolae for potential vesicular trafficking into and across the cell. Unlike passive transvascular delivery, which requires large doses to generate sizeable concentration gradients across the blood-tissue interface to drive tissue entry, caveolae constitute a pumping system in ECs that can bud to form dynamic discrete vesicles ^20, 21, 22^ that transcytose select bound probes, even at low doses and against a sizeable concentration gradient ^23, 24, 25, 26, 27, 28^. One such probe is mAPP2, a lung caveolae-targeting antibody (CTA) generated to aminopeptidase P2 (APP2) which is highly clustered in lung EC caveolae *in vivo* ^23, 27, 29^. Caveolae effectively concentrate mAPP2 in lungs quite specifically by extracting most of it from the blood and pumping it across the EC barrier to flood the lung interstitium within minutes of iv injection ^23, 26, 27, 28, 30^. Little mAPP2 is left in the blood or other organs. No other antibody or probe to date has demonstrated similar precise, robust and rapid targeting and tissue penetration.

Most NPs exhibit a natural tropism for the RES that can approach the CPS in blood extraction speed and efficiency, albeit into multiple tissues of the RES. Whether caveolae can also pump iv injected CTA-conjugated NPs as effectively to concentrate them only inside a single tissue is unknown and rendered uncertain because of the small size of caveolae and the potentially overpowering ability of the RES to clear NPs from the blood within minutes. Caveolae form flask-shaped invaginations with ∼70 nm outer diameter bulbs; their necks provide 20-30 nm circular openings at the vessel wall that face outwardly and perpendicular to the blood flow ^27^. Hence, both size and hydrodynamic forces as well as an extensive EC glycocalyx expected *in vivo* ^31^ may significantly hinder NP entry into caveolae from the circulating blood. So far all mAPP2 derivatives (Fab, scFv, bispecific) showing lung targeting and penetration have been near IgG size, readily <15nm in hydrodynamic diameter (HDD) ^23, 26, 30, 32^. The extent of pumping of CTA-conjugated NPs remains unknown and requires detailed quantification *in vivo*. Composition and size of the NPs are arguably the two main contributors to reduced target-tissue uptake.

Here, we begin to study these key issues and possible nano-pumping via caveolae by coupling a variety of NPs to mAPP2 to visualize and quantify tissue targeting after iv injection. We investigate the extent and specificity of caveolae-targeting as a function of NP’s size. We characterize trafficking limitations and estimate optimal diameter range of functionalized NPs for efficient caveolae-targeted delivery. These parameters will be critical in the design of NP carriers for potential diagnostic imaging and/or therapeutic applications using the caveolae transcytosis pathway. The key questions answered here include: will the RES or the CPS in the lung prevail for NP uptake, will the CTA be effective in retargeting NPs or will the NPs end up retargeting the antibody to the RES, and to what degree, if any, will NP size and type change the outcome? Ultimately, we discover that CPS targeting can enhance NP delivery to lung by orders of magnitude and that tissue-precise, active nanodelivery across restrictive continuous endothelium is indeed possible by redirecting NPs to the CPS, a new, robust and active pathway to get beyond the current inadequate paradigm of passive transvascular delivery of NPs.

## Retargeting dendrimers to test lung CPS vs RES scavenging

Our general working hypothesis is that the CPS in microvascular endothelia can compete with the RES for rapid and robust NP extraction from the circulating blood. Specifically, when linked to NPs, the lung EC CTA mAPP2 will redirect their biodistribution after iv injection from the liver and spleen towards precision lung uptake. We started to test this hypothesis qualitatively using very small NPs to constrain minimally caveolae entry and APP2 binding, thereby increasing the probability of lung targeting and penetration of the EC barrier.

We first iv injected radiolabeled G5 PAMAM dendrimers and G4 PAMAM dendrons (HDD 5 and 4 nm, respectively) conjugated to mAPP2 Fab as depicted in Suppl. Fig. **1**. Planar γ-scintigraphy and SPECT-CT whole body imaging as well as biodistribution analysis of excised organs all supported the hypothesis by showing the CTA strikingly altered the biodistribution profile of the NP towards lung uptake and away from RES uptake (Fig. **1A-L**). Using SPECT-CT imaging, we observed rapid and robust lung-specific accumulation of mAPP2-G5 dendrimers (Fig. **1B**). A very distinct lung shape was evident with a negative signal in the mediastinum (negative ghost cardiac image), consistent with little radioactivity remaining in the blood. This high intensity lung signal occurred primarily within 30 min and persisted even a week after iv injection (Fig. **1C**). In contrast, the non-targeted G5 dendrimers rapidly and robustly accumulated predominantly in the liver and spleen, also mostly within 30 min (Fig. **1A** and **1D**).

**Figure 1.**
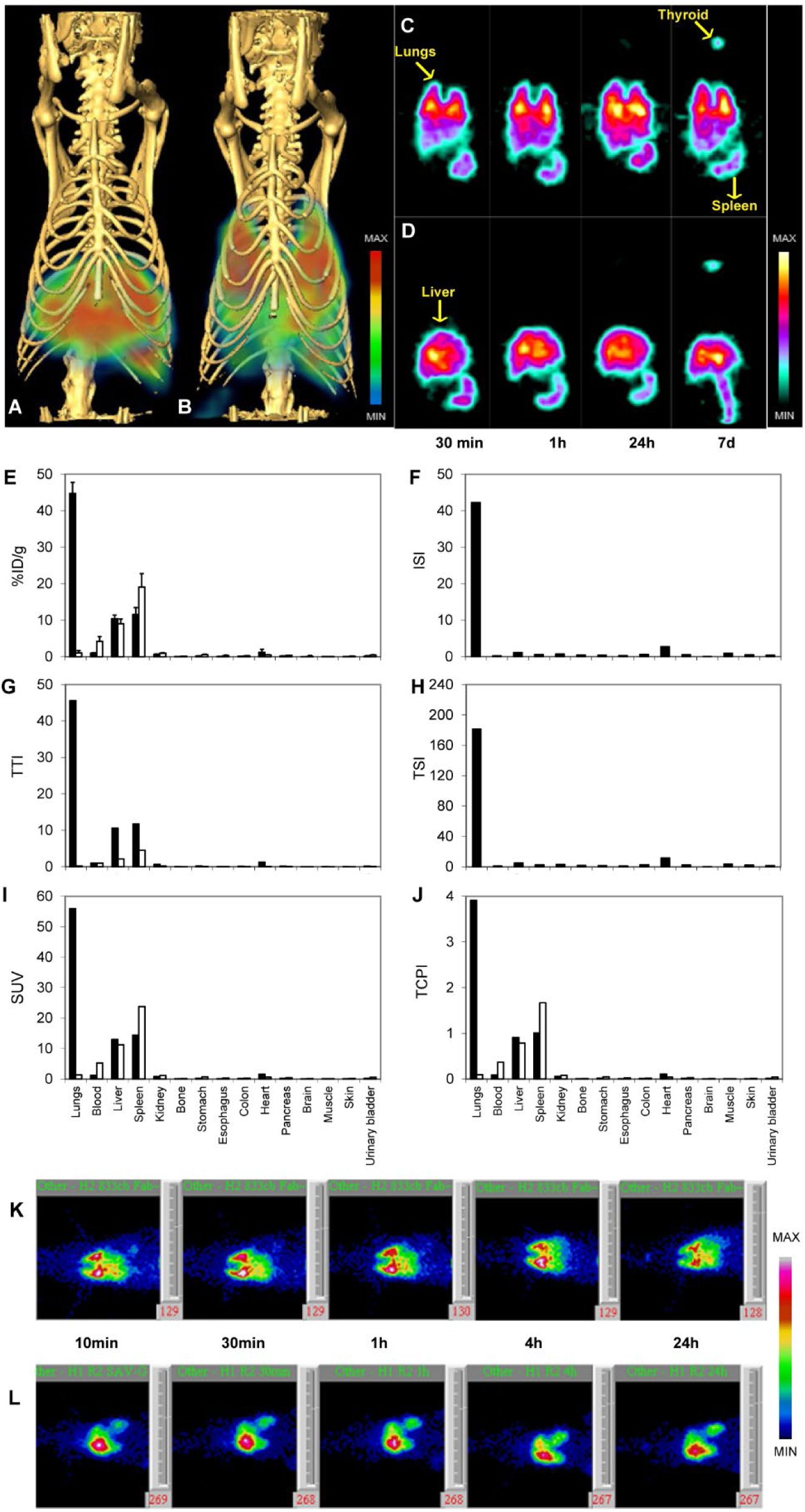
Lung-specific retargeting, biodistribution analysis and targeting indices of APP2-targeted PAMAM dendritic nanoparticles. 3D-images of CT-SPECT fusion of volumetric SPECT texture and skeletal CT were acquired at 1 h after iv injection of (**A**) nontargeted or (**B**) mAPP2 Fab conjugated ^125^I-labeled PAMAM G5 dendrimers. Tomographic SPECT images of APP2-targeted (**C**) PAMAM G5 dendrimers and (**D**) untargeted dendrimers are shown at the indicated time after administration. Tissue uptake (%ID/g), (**E**), ISI (**F**), TTI (**G**), TSI (**H**), SUV (**I**), and TCPI (**J**) were determined in indicated organs and blood at 1 h after iv injection of APP2-Fab targeted or control nonspecific Fab conjugated ^125^I-labeled PAMAM G5 dendrimers. N=5 for all experimental groups. Data are expressed as means with corresponding standard deviation (SD). See Methods for statistical details. Planar γ-scintigraphic imaging of APP2-Fab retargeted (**K**) or nontargeted (**L**) ^125^I-labeled G4 dendrons at the indicated time after iv injection.

Biodistribution analysis of excised organs confirmed this very effective lung retargeting of the NPs. Lung uptake of mAPP2-G5 dendrimers reached 45% ID/g at 1 hr, compared to 1.1% ID/g for untargeted G5 dendrimers, in this case linked to a nontargeting control p27 Fab (Fig. **1E**). These two values as a ratio yield an impressive immunospecificity index (ISI) of 42, far greater than detected for any other organ (Fig. **1F**). Without further NP cloaking to block rapid RES uptake, caveolae pumping was still able to boost NP delivery to lungs impressively by >40-fold despite direct competition from the RES.

As expected, both the RES and CPS cleared the blood of radioactivity rapidly and efficiently (Fig. **1C-E**). The blood levels were very low at 1 h (Fig. **1E**). The mAPP2-G5 dendrimers exhibited the lowest blood level at 1% ID/g, probably benefiting from dual extraction. The tissue to blood ratio (tissue targeting index (TTI)) was highest for lung at 46. This readily confirmed lung targeting was achieved, but some liver and spleen targeting was still evident, in part boosted by the ultra-low blood concentration (Fig. **1G**). The tissue specificity index (TSI) might be a more informative gage because it normalizes the TTI by the TTI of the untargeted NP, thereby comparing the two directly to show how the tissue selectivity of the probe has changed. Fig. **1H** shows clear lung specificity; the impressive lung TSI above 180 quantified how much active delivery by caveolae can exceed the customary passive delivery paradigm for NPs even in the face of stiff competition from RES scavenging.

We also analyzed the NP targeting, retention efficiency and concentrating power of the caveolae transport pathway using two additional measures: the specific uptake value (**SUV**) and the tissue concentrating power index (**TCPI**). SUV quantifies the improvement caveolae targeting brings to NP delivery compared to an ideal probe that penetrates all tissue compartments homogenously (Fig. **1I**). Values above 1 suggest preferential targeting. TCPI is an even more demanding index because it normalizes the tissue uptake relative to the peak blood concentration which drives passive transvascular delivery and therefore rarely is reached inside organs with restrictive EC barriers (Fig. **1J**). Values are expected to be and usually are small fractions and approaching 1 or more requires more than just passive delivery. The EC CPS exhibited a robust ability to concentrate dendrimers in the lungs with very high SUV and TCPI values for NPs of 56 and 3.9, respectively (Fig. **1I** and **1J**). Ultimately, caveolae can concentrate NPs inside lung to levels 4-fold greater than their peak level in the blood (100% ID immediately after iv injection). These results are unprecedented for NPs and even exceed the concentrating power of the RES which still impressively produced a TCPI near 1 for the liver and spleen. The RES was even more impressive in concentrating the control Fab conjugated NPs with the spleen SUV and TCPI reaching 24 and 1.7 respectively. Both CPS and RES are superb at concentrating these small NPs and the CPS can redirect NPs to the lung.

Similarly, imaging by γ-scintigraphy of mAPP2-G4 dendrons also showed rapid lung-specific targeting at 10 min (Fig. **1K**) unlike untargeted G4 dendron constructs which appeared to accumulate primarily in the liver and spleen (Fig. **1L**). ROI quantification (**Table 1**) confirmed delivery of mAPP2-G4 dendrons into the lungs quickly exceeded liver uptake by 4.0-fold at 10 min with a maximum of 4.5-fold at 30 min. CTA retargeting was striking with 15-fold more total lung uptake than the untargeted G4 dendrimer. Also, liver uptake was markedly reduced by >5-fold. Retargeting by mAPP2 conjugation enhanced the lung to liver uptake ratio by 64-fold at 10 min, well before deiodination takes place in the liver.

**Table 1.** Time-dependent uptake of APP2-targeted PAMAM G4 dendrons. The uptake of mAPP2-Fab retargeted and nontargeted ^125^I-labeled G4 dendrons in lung and liver tissues was quantified from planar g-scintigraphic acquisitions using LumaGEM_®_ software at indicated time points (n=4).

|  |  | %ID in the ROI |  |  |  |  |
| --- | --- | --- | --- | --- | --- | --- |
|  | Organ | 10 min | 30 min | 1 h | 4 h | 24 h |
| APP2Fab~SAV-G4 - <sup>125</sup>I | Lung | 55.9 | 54.4 | 50.0 | 45.8 | 35.4 |
|  | Liver | 14.1 | 12.1 | 11.6 | 10.9 | 10.7 |
| Untargeted G4 dendron- <sup>125</sup>I | Lung | 3.8 | 4.7 | 3.4 | 3.8 | 2.9 |
|  | Liver | 61.3 | 69.4 | 64.6 | 57.5 | 56.2 |

## Characterizing Au NP immunoconjugates

Although the above data support our hypothesis clearly, there are expected limitations in the size of NPs that can enter caveolae and bind APP2 to then be redirected into the lung. To address size effects, we chose hard, inflexible and spherical colloidal gold NPs (GNPs) commercially available with well-defined, narrow size distributions. As depicted in supplemental Fig. 2, GNPs of 10 nm, 20 nm, and 40 nm diameter cores were functionalized with PEG12-carboxylic acid and conjugated directly with one of two whole IgGs: mAPP2 or mAPP2X which has 2 amino acids mutated to abolish any APP2 binding and thus represents an almost physicochemically identical and ideal control. The final hydrodynamic diameter (HDD) of both immunoconjugated GNPs as measured by dynamic light scattering (DLS) was equivalent for each Au NP size: GNP10 at 23 nm, GNP20 at 33 nm and GNP40 at 64 nm. These NP complexes were physicochemically and biologically characterized (see Table 2, Fig. 2 and Suppl. Fig. 2). The results demonstrate successful antibody conjugation to the GNPs with desired range of polydispersity index and zeta potential for in vivo delivery and APP2 specificity of mAPP2 covalently bound to the NPs (Fig. **2B** & Table **2**). The mutated version mAPP2X, did not bind APP2 (Fig. **2C**) by design and their respective GNP conjugates had no apparent affinity towards APP2 (Suppl. Fig. 3). Prior to their *in vivo* application, radio-immunonanoconjugates (RINCs) were prepared by directly labeling of antibodies (mAPP2 or mAPP2X) with ^125^I and further conjugation of radiolabeled antibodies with the GNPs, resulting in radiochemical yields of 70-85%. The average number of antibodies per GNPs as estimated from molar particle concentrations and specific activities were 1, 4 & 36 (Table **2**) for RINCs with 10, 20 and 40 nm core diameters, respectively. Additionally, RINCs were stable in human serum after 24 h at 37°C.

**Figure 2.**
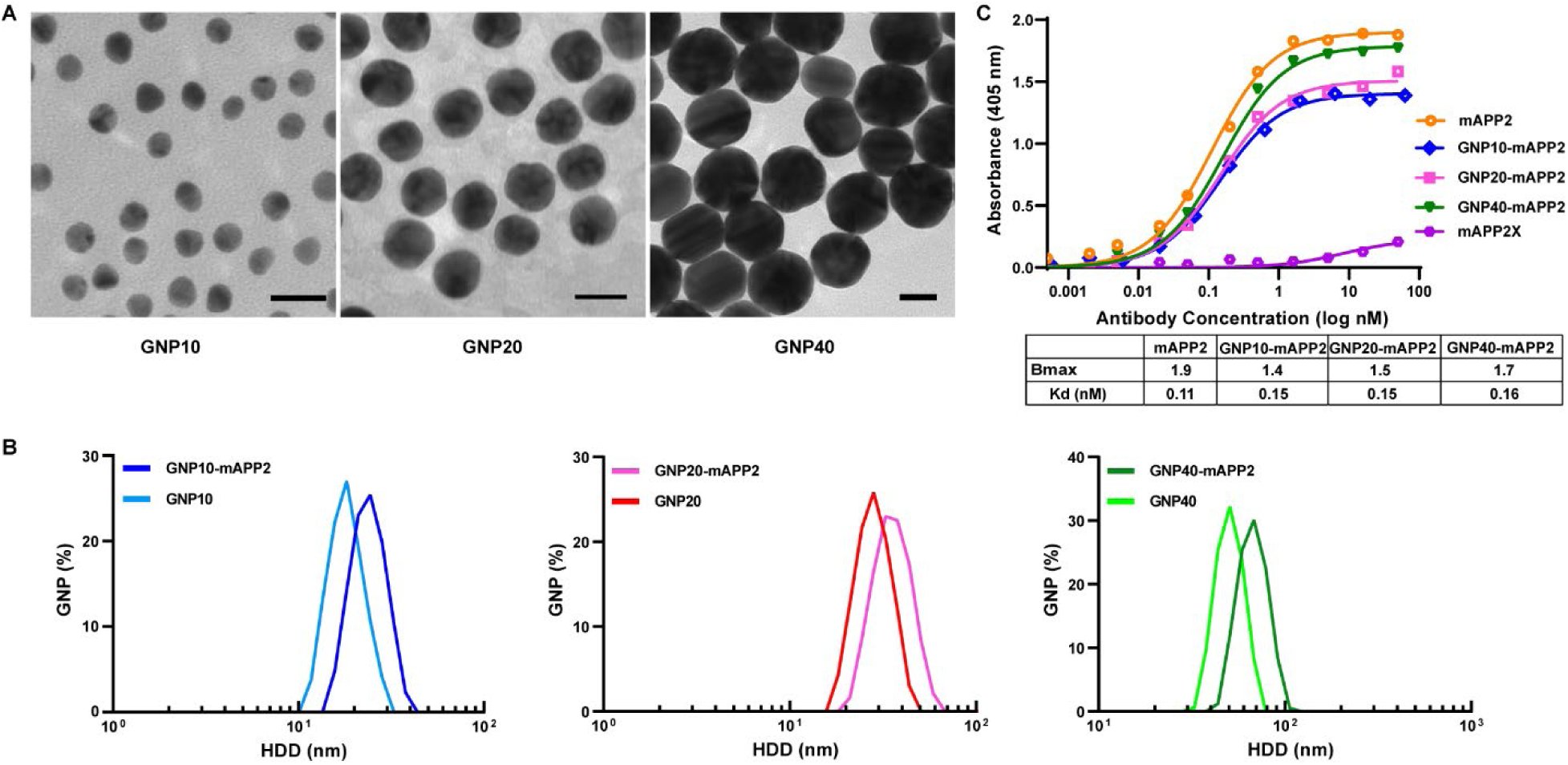
Characterization of GNPs (10, 20 and 40 nm) and respective GNP immunoconjugates. (**A**) Representative transmission electron micrographs (TEM) of PEG_12_-carboxylic acid functionalized GNPs (scale bar = 20 nm) showing core diameter for GNP10 (10.8±0.9 nm), GNP20 (19.1±2.4 nm) and GNP40 (40±4 nm). (**B**) Dynamic light scattering (DLS) of GNPs with or without conjugated antibody (mAPP2/mAPP2X) showing average hydrodynamic diameter (HDD) of GNP10 (17.8 nm), GNP10-mAPP2 (23.4 nm), GNP20 (26.4 nm), GNP20-mAPP2 (33.3 nm), GNP40 (46.5 nm) and GNP40-mAPP2 (63.6 nm). (**C**) Apparent affinities (Kd) of antibodies or GNP conjugated antibodies measured by ELISA. See methods for additional details.

**Table 2.** Physiochemical characterization of immunonanoconjugates in comparison to unconjugated GNPs and antibodies.

|  | mAPP2/X | GNP10 | GNP10-mAPP2/X | GNP20 | GNP20-mAPP2/X | GNP40 | GNP40-mAPP2/X |
| --- | --- | --- | --- | --- | --- | --- | --- |
| **Core Diameter (nm)** |  | 10.8±0.9 | - | 19.1±2.4 | - | 40±4 | - |
| **Hydrodynamic Diameter (nm)** | 11±0.4 | 17.8±4.8 | 23.4±5.2 | 26.5±6.5 | 33.3±7.7 | 46.5±8.7 | 63.6±11 |
| **Polydispersity Index** |  | 0.14 | 0.18 | 0.13 | 0.2 | 0.12 | 0.16 |
| **Zeta Potential (mv)** |  | -35 | -11 | -46 | -9.8 | -38 | -7.9 |
| **Molar Particle Concentration (nM)** |  | 66 | 46 | 16 | 12 | 1.9 | 1.4 |
| **Antibody/Particle** |  |  | 1 |  | 4 |  | 36 |
| **Serum stability** | 97% |  | 97% |  | 96% |  | 96% |

## Imaging NP size effects on lung vs RES targeting

To evaluate the targeting potential and uptake kinetics, RINCs were administered iv into female SD rats. Because both lung caveolae pumping and RES scavenging can mediate bulk extraction from blood into tissue in just a few minutes, we performed rapid acquisition, planar whole-body imaging on rats starting minutes after iv injection of the RINCs. Consistent with the expected rapid uptakes and expected *in vivo* release of ^125^I from the probes, the planar ***γ***-scintigraphic images (Fig. **3A**) were the most robust, distinguished and informative at the early time points (10 min and 1 h). They showed ample, rapid and distinct lung uptake/targeting that clearly diminished with increasing NP size. The antibody alone gave a very strong, unequivocal lung image with a negative cardiac ghost image even evident at 10 min (see arrowhead); the lung signal was much greater than any blood-engorged organ signal like liver or heart, consistent with rapid and extensive blood extraction. The lung image became less distinct and robust with increase in GNP size. The GNP10 conjugate lung image was not attenuated much but the liver image became more apparent. These effects were a bit more pronounced with GNP20 and peaked with the GNP40 conjugate yielding a very distinct and dominating liver image, even at 10 min with low lung uptake that can nevertheless still be detected. In many ways, the images reversed from rapid precision lung uptake to rapid precision liver uptake once the Au core diameter reached 40nm and the overall nanoconjugate exceeded 60 nm in HDD.

**Figure 3.**
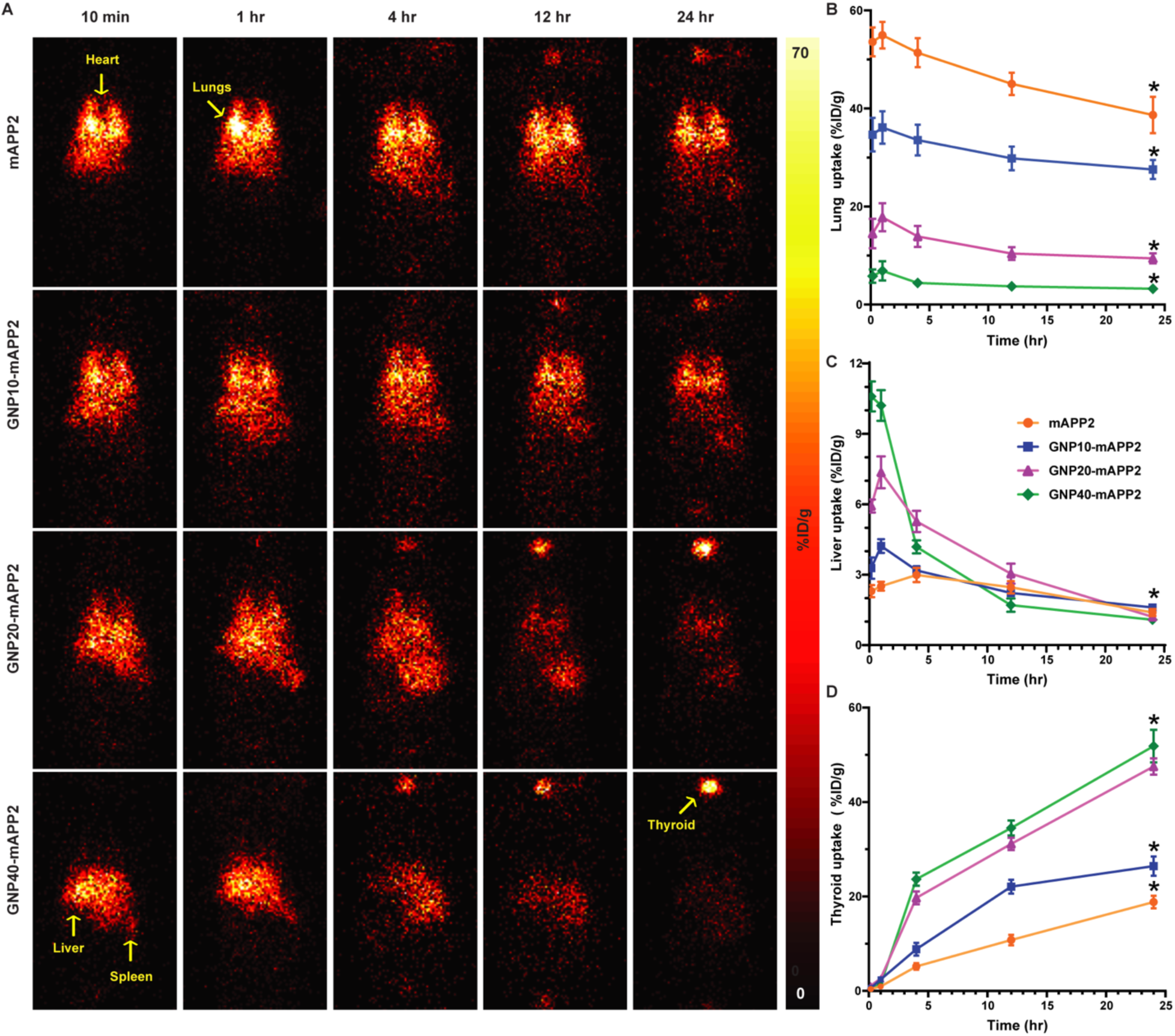
*In vivo* tracking of lung EC caveolae-targeted GNPs with increasing core diameters. (**A**) Serial planar ***γ***-scintigraphic images captured at 10 min, 1 hr, 4 hr, 12 hr & 24 hr time points after iv injection of either radioimmunoconjugate (mAPP2-^125^I) alone or conjugated with GNPs of indicated core diameters (GNP10/20/40-mAPP2-^125^I). (**B**) Image-based quantitation of lung uptake over time for caveolae-targeted GNPs with indicated core diameters. (**C**) Image-based quantitation of liver uptake over time for caveolae-targeted GNPs with indicated core diameters. (**D**) Image-based quantitation of thyroid uptake over time for caveolae-targeted GNPs with indicated core diameters. *Uptake was significantly lower for lungs and liver, and higher for thyroid, at 24 hr compared to the 10 min and 1 hr time points (Bonferroni, α=0.05).

Looking at the collage of images as an ensemble readily shows radioactivity accumulated primarily in lung, RES and thyroid. We performed ROI analysis to quantify this uptake for each probe and organ. The lung reached maximal uptake within minutes (10 min and 1 h were not statistically distinct) for all the probes (Fig. **3B**). The liver uptake was also rapid but the GNP10 and GNP20 immunoconjugates showed ∼20% more signal at 1 h than 10 min (Fig. **3C**). The lung signal decreased modestly (5-25%) over time independent of NP size whereas the liver signal diminished much more rapidly and extensively in accord with the degree of original uptake and NP size. These signal attenuations are customary from *in vivo* dehalogenation processes leading to uncoupling the ^125^I label from the antibody as has been well documented to occur in the RES and blood ^33, 34^. As expected, the thyroid with its specific iodine pumping mechanism extracted and accumulated some of the freed ^125^I (Fig. **3C**). The images and quantification show no thyroid signal at early time points (indicating little free unconjugated ^125^I at injection). Noticeably, the thyroid becomes apparent first for the larger nanoconjugates, clearly at 4 h maybe sooner. The ROI analysis shows impressive thyroid accumulation especially pronounced for the 20 and 40 nm gold core conjugates, likely from greater RES uptake and dehalogenation.

Motivated by the above results, we focused future experiments on the 1 h post-injection time point which appears equitable to both RES and caveolae pumping while obviating complications from the ^125^I uncoupling. We utilized SPECT-CT along with planar imaging to visualize biodistribution of APP2 and lung caveolae targeting ability of RINCs (GNP10/20/40-mAPP2-^125^I) in comparison to respective nontargeted RINCs (GNP10/20/40-mAPP2X-^125^I). As can be seen from the SPECT-CT images (Fig. **4A** & **4B**), specifically the maximum intensity projection (MIP) images confirmed the planar imaging results (Fig. **3A**) while revealing the striking difference between the paired equivalent probes. mAPP2X lost the ability of mAPP2 to target lung and produced a customary nonspecific antibody signal circulating in the body, namely poor intensity in any single organ but most readily apparent in blood-rich organs like heart, liver and lung. Unlike mAPP2, no cardiac ghost image was observed here because nearly all injected mAPP2X resides in the blood. The differences between the NP pairs could be just as striking with excellent lung precision targeting being lost when using mAPP2X. Both GNP10 and GNP20 appeared driven into lung when conjugated to mAPP2. But with mAPP2X the NP’s natural liver tropism took over the targeting. This was especially pronounced for GNP40 which pushed both antibodies into the liver to generate robust liver uptake.

**Figure 4.**
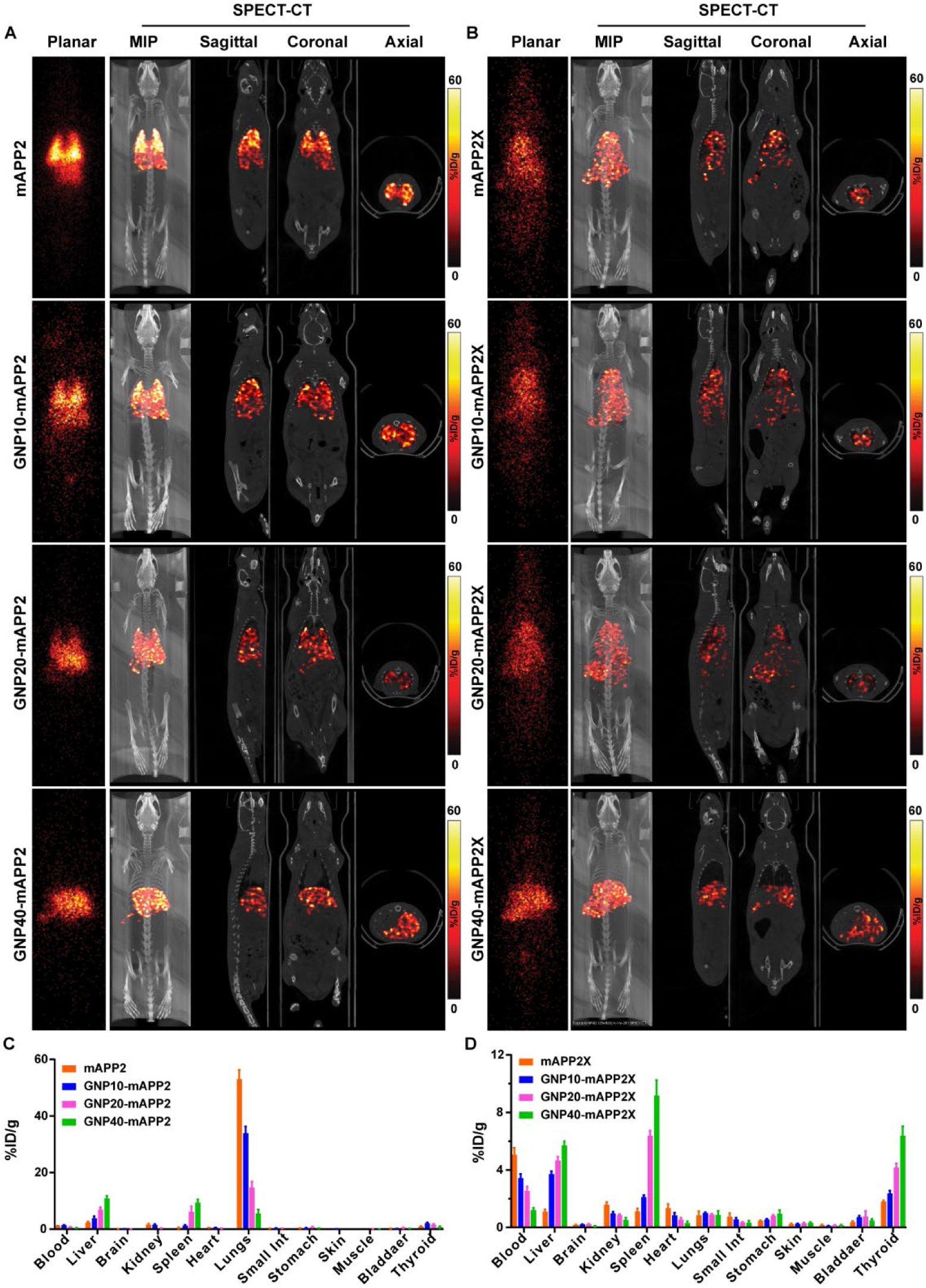
*In vivo* planar *γ*-scintigraphic, SPECT-CT imaging and biodistribution of targeted and non-targeted immunoconjugates with or without GNPs of different core diameters after 1 hr injection in normal female SD rats. (**A**) Representative planar *γ*-scintigraphic and SPECT-CT (MIP, sectional) images captured at 1 hr after iv injection of either targeted radioimmunoconjugate (mAPP2-^125^I) alone or conjugated with GNPs of indicated core diameters (GNP10/20/40-mAPP2-^125^I). (**B**) Representative planar *γ*-scintigraphic and SPECT-CT (MIP, sectional) images captured at 1hr after iv injection of either non-targeted radioimmunoconjugate (mAPP2X-^125^I) alone or conjugated with GNPs of indicated core diameters (GNP10/20/40-mAPP2X-^125^I). (**C**) Biodistribution of GNP-mAPP2-^125^I in comparison to mAPP2-^125^I at 1 h after injection in normal female SD rats (130-140 g). (**D**) Biodistribution of GNP-mAPP2X-^125^I in comparison to mAPP2X-^125^I at 1 h after injection in normal female SD rats (150-160 g). While separate animals (n=1) were used for SPECT-CT imaging, animals subjected to 1 hr planar imaging (n=3) were also utilized for quantitative biodistribution study. Data expressed as mean ± SD.

## Biodistribution analysis and targeting indexes quantify size effects

The results from the planar and SPECT-CT imaging were further corroborated by quantitative biodistribution studies (Fig. **4C** & **4D**). Targeted RINCs, specifically GNP20 and GNP10, showed significantly higher uptake in lungs (14 - 34% ID/g) compared to their nontargeted equivalents showing poor lung uptakes (<1% ID/g) and a different biodistribution profile dominated by signal in blood, liver, spleen and thyroid. Again, GNP40, both targeted and non-targeted RINCs, had maximum RES accumulation reaching ∼10% ID/g.

To evaluate the tissue specificity and extent of mAPP2-targeted delivery of GNPs as a function of their size (core diameter or HDD), we used the biodistribution data to calculate different targeting indices. For the lungs of rats injected with targeted RINCs (mAPP2-^125^I or GNP-mAPP2-^125^I), ISI values for lung exceeded all other tissues for each GNP core diameter studied. Fig. **5A** shows mAPP2 increases lung uptake significantly for all constructs relative to the X controls by >60 times, >30 for GNP10, 20 for GNP20, and even 6 for GNP40 constructs. Also note in Fig. **5B** that the NP tropism for RES robustly retargeted the mAPP2X towards the spleen and liver with as much as 8-fold increase to spleen for the GNP40 construct. Even GNP10 increased liver uptake of mAPP2X by >3-fold. To quantify the accumulation of mAPP2-GNPs in specific tissues relative to their remaining levels in blood circulation, we calculated TTIs. For lungs, TTIs were highest for mAPP2-GNP10 and mAPP2-GNP20 (Fig. **5C**). However, with GNP40, TTIs of liver and spleen (Fig. **5C** **& 5D**) were higher for both targeted and non-targeted immunoconjugates in comparison to TTIs of lungs. These values indicate that mAPP2-GNP complexes with HDD ≤ 35 nm (unlike the larger 64 nm NPs) are more efficiently removed from the blood and transported into lung tissue than into off-target RES tissues.

**Figure 5.**
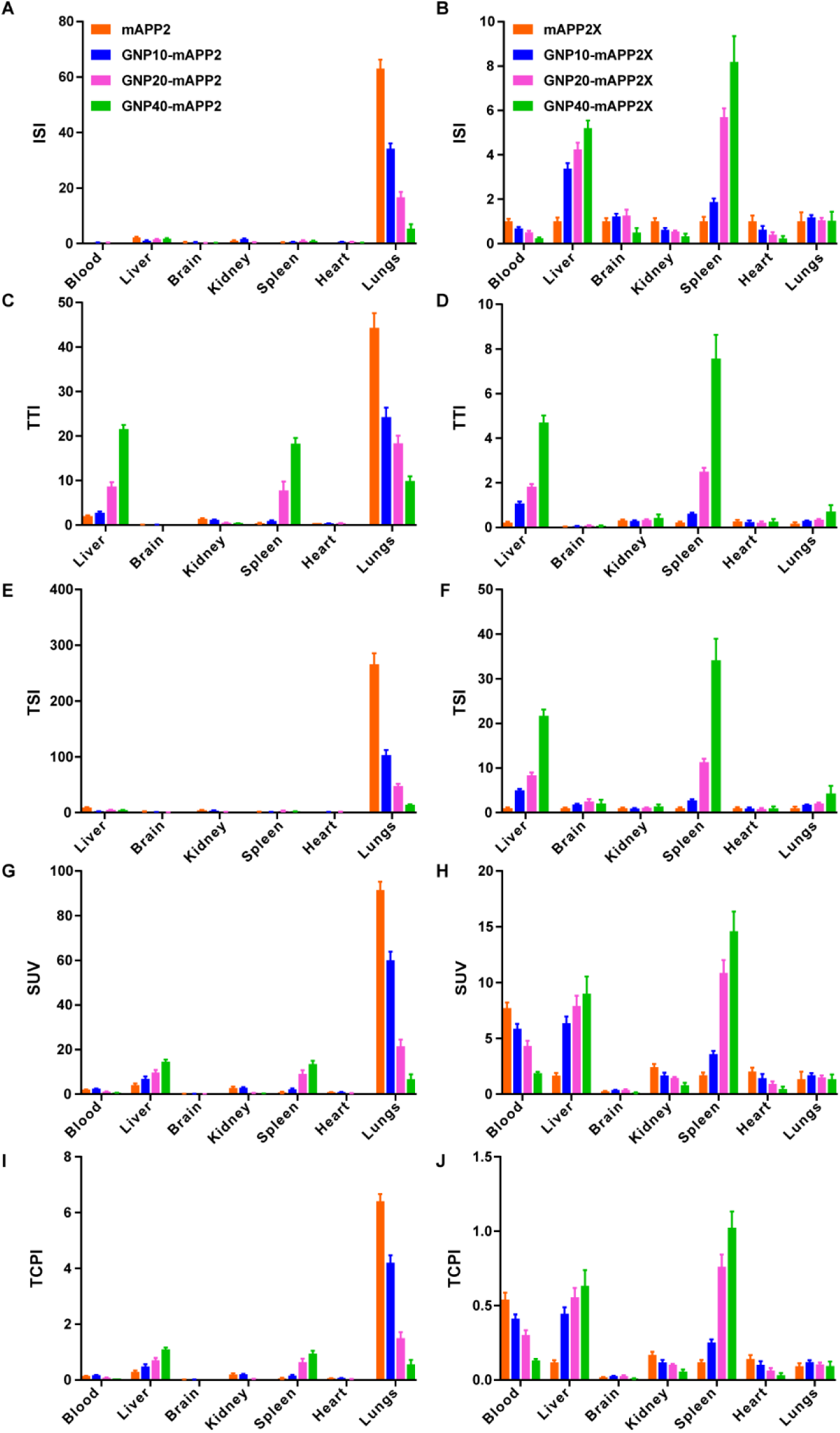
Targeting indices generated using biodistribution data for evaluation of target specificity, delivery and pumping of caveolae-targeted GNPs as a function of their size. Following targeting indices were calculated for GNP-mAPP2-^125^I and GNP-mAPP2X-^125^I: **(A)** Immunospecificity index (**ISI**) as a ratio of tissue uptake (%ID/g) of targeted NPs (GNP-mAPP2-^125^I) to the tissue uptake of respective control non-targeted NPs (GNP-mAPP2X-^125^I). (**B**) For ISI of GNP-mAPP2X-^125^I, the uptake was normalized to uptake of negative control alone (mAPP2X-^125^I), (**C&D**) Tissue targeting index (**TTI**) as a ratio of tissue to blood uptake, (**E**) Tissue specificity index (**TSI**) as a ratio of TTI of targeted NPs versus TTI of control non-targeted NPs. (**F**) For TSI of GNP-mAPP2X-^125^I, TTI was normalized to TTI of negative control alone (mAPP2X-^125^I), **(G&H)** Specific uptake value (**SUV**) was calculated as tissue uptake normalized to the uptake that would be reached after homogeneous penetration throughout all tissues, and (**I&J**) Tissue concentrating power index (**TCPI**) was estimated as a ratio of uptake in each tissue to the peak blood level. N=3 for all experimental groups. Data are expressed as mean ± SD.

To compare further tissue targeting properties of caveolae-targeted GNPs to their non-targeted counterparts, we calculated the TSI for each organ which revealed again excellent redirected lung specificity. For lungs, the TSIs of caveolae-targeted GNPs exceeded 100 for GNP10, 45 for GNP20, and 15 for GNP40, and these values were well above TSI estimates for other off-target-tissues (< 6) (Fig. **5E**). Conversely, the TSIs of both liver and spleen were higher for nontargeted GNPs, which increased with NPs size (Fig. **5F**). These results provide compelling evidence that caveolae-targeting greatly improves delivery of NPs to target tissue compared to passive transvascular delivery mechanisms.

Both SUV and TCPI confirmed the power of CPS to concentrate each NP size inside lungs (Fig. **5G** **& 5I**). The SUV was significantly greater in the lungs than other tissues, particularly for targeted RINCs with 10 nm (60), 20 nm (20), and 40 nm (8) cores. The SUV of the nanocomplexes with HDD of 64 nm was the highest in the liver and spleen ranging from 9-17 regardless of which antibody was conjugated to the NPs (Fig. **5G** **& 5H**). The TCPI was greater than 1 only in lung and only for the caveolae targeted GNP10 (4.2) and GNP20 (1.5). These data demonstrate lung endothelial CPS exhibit a significant and size-selective concentrating power for mAPP2/caveolae-targeted NPs with HDD ≤ 35 nm. Ultimately the lung CPS can concentrate NPs inside lungs to levels 4-fold greater than their peak level in the blood. These results are unprecedented and even exceed the concentrating power of the RES which still impressively produced a TCPI of 1 for the 64 nm complexes (Fig. **5I** & **5J**).

To prove that the antibody and the GNP were distributed together, we also used inductively coupled plasma mass spectrometry (ICP-MS) to measure the actual amount of gold in 5 key tissues 1 h after iv injection of GNP10 and GNP40 immunoconjugates. These results (Fig. **6A**) were very similar to the results obtained using radiolabeling (Fig. **4C** & **4D**). The Au NPs were clearly redistributed to lung by mAPP2 conjugation, yielding 36% ID/g for GNP10 and 6% ID/g for GNP40. Normally these NPs target the RES robustly with little lung uptake. Regardless of detection system used, Fig. **6B** & **6C** show that with mAPP2 conjugation the delivery ratio of lung to liver or spleen goes from well below 1 to nearly 1 for GNP40 to 100+ for GNP10 in the spleen.

**Figure 6.**
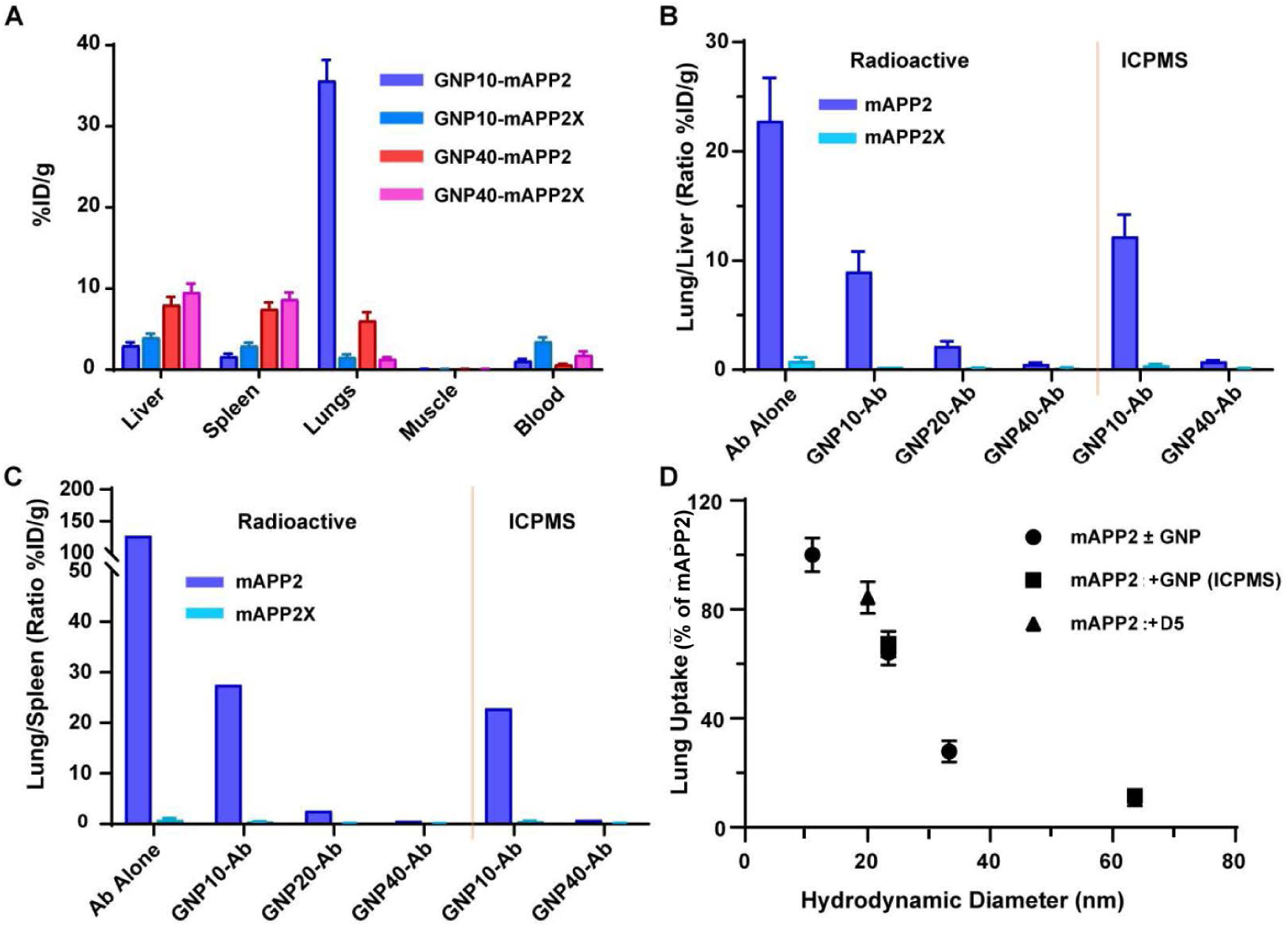
(**A**) Biodistribution of gold uptake as determined by ICP-MS performed on organs of interest harvested from animals injected iv with GNP10-mAPP2-^125^I & GNP40-mAPP2-^125^I and compared with corresponding controls (GNP10-mAPP2X-^125^I & GNP40-mAPP2X-^125^I). N=3 for each animal group. Partition coefficient in terms of (**B**) Lung/Liver uptake, (**C**) Lung/Spleen uptake for both radioactivity and gold content as determined by ex vivo biodistribution studies. (**D**) Relative lung uptake for all nanoconjugates with varying HDDs (dendritic (D5) NPs and GNPs) obtained in terms of radioactivity and gold content by ex vivo biodistribution.

Lastly, Fig. **6D** shows the partitioning of all mAPP2 nanoconjugates into lung tissue as a function of size. This partition function clearly shows the effects of HDD on caveolae pumping. Although significant lung uptakes were evident for all HDDs tested, it appears from the curve tendency that sizes beyond 100 nm will likely be least fruitful and that sizes below 25 nm are best with the least effect on uptake. Also, it implies that probes below the size of an antibody may possibly show even more pumping. Better cloaking of the NPs to slow down or hopefully obviate RES scavenging will likely enhance lung caveolae pumping even further.

## New data in context of past reports

High-precision tissue delivery is critical for NPs to be useful in the imaging and therapy of many diseases. We and many others before us, several decades ago before the term NP was even invented, used ferritin and protein-coated colloidal gold particles to reveal a role for plasmalemmal vesicles (now called caveolae) in their trafficking into ECs and across restrictive EC barriers in various non-RES tissues ^9, 35, 36, 37, 38, 39^. These discrete, large and electron dense probes were easy to image by electron microscopy (EM); they were too big for junctional transport and no intercellular transport was detected. But rather, entry of multiple NPs into apparently single caveolae at the luminal EC surface was detected followed later by exit of similarly numbered NPs from abluminal caveolae into the tissue interstitium. Cumulatively, these EM studies and their extensions to cultured cells portrayed caveolae not as obligate sessile structures but rather as facultative, dynamic traffickers of these probes into many cell types (endocytosis) and across ECs (transcytosis) ^9, 35, 36, 37, 38, 39^. Both transport processes appeared rather slow and inefficient.

More recently, we performed EM on lung tissue after pulmonary arterial perfusion of mAPP2 coupled to 10-15 nm Au NP *in situ* to provide static, high magnification images that confirmed specific lung EC surface binding of mAPP2 occurred inside caveolae^23, 27^. Only when coupled to mAPP2 and not isotype matched IgG, the Au NPs were concentrated inside caveolae before being transcytosed into the lung interstitium. Transcytosis for all of these electron-dense NPs was examined in the past primarily under artificial *in situ* perfusion conditions, not iv injection. Under these less than ideal physiological and hemodynamic conditions, they appeared to require high intravascular concentrations to achieve caveolae entry and transcytosis that produced interstitial concentrations well below the levels observed originally inside the blood vessels. Although these studies clearly showed that NP can passively enter caveolae which can deliver them, bound or not, across the endothelium, the pathway appeared quite limited, causing some to question their dynamic nature and any realistic physiological role in extravasation^40, 41, 42, 43^. Subsequent studies cemented a dynamic function once purification and proteomic mapping of EC caveolae from lung tissue revealed that caveolae, like clathrin-coated vesicles, shared related docking and fusion proteins that used high energy GTP/ATP hydrolysis to drive trafficking and that could be inhibited by mutation or pharmacological means to prevent transport^20, 21, 22, 28, 29, 44, 45, 46,47^.

Here our experiments reveal and quantify for the first time the exceptional efficiency of the CPS for NP delivery, in particular immunotargeting caveolae to redirect NPs towards the lung and away from the RES. Our findings demonstrate the rather striking, potentially critical discovery that caveolae targeting can actually concentrate NPs rapidly in a single tissue with unprecedented precision and at extraordinary levels well beyond the highest blood concentration after iv injection. Therefore, CPS targeting is an effective, systemic delivery strategy for NPs and their cargo. Here the NP cargo was used for imaging biodistribution and constituted the radionuclide bound either directly to the dendrimers and dendrons or indirectly via the antibody conjugated to the Au NP.

Our results quantified in multiple ways show lung NP delivery can be boosted by one to two orders of magnitude over controls including mAPP2X mutated in just 2 amino acids to ablate APP2 binding. These data contradict a mechanism mediated solely by passive transport through sessile luminal and abluminal caveolae linked to form transendothelial channels for passive diffusive and convective transport. But rather they support active and discrete vesicular transcytosis of the bound NP. The control untargeted mAPP2X NPs, which should utilize any passive pathway as effectively or more than targeted mAPP2 NPs, exhibited much less extravasation. Binding to APP2 at the walls of static caveolae channels should slow, impede and ultimately limit not facilitate passive filtration. A simple analogy is size exclusion liquid chromatography based on filtration through beads with pores having inert or interactive surfaces; binding of the same size probe in the channel retards its flux. On the contrary, an active dynamic transcytotic mechanism would readily benefit from specific binding to concentrate more of the probe inside each discrete vesicular trafficking unit and thereby substantially boost NP transcytosis of mAPP2 conjugates over mAPP2X conjugates relegated only to fluid phase uptake and other shared passive conduits. Thus, these new data provide further evidence of i) the dynamic not sessile nature of caveolae in endothelium *in vivo* and ii) the robustness and ample capacity of the CPS for precision delivery, targeting, and imaging.

## Size matters for CPS to overcome RES

We examined 6 distinct targeting indices, all of which clearly indicate that caveolae targeting of dendrimeric and Au NPs achieves precision NP delivery to lung at levels that not only significantly surpass their passively delivered, nontargeted counterparts but also can even greatly exceed the highest RES uptakes. The partition function curve (Fig. **6D**) probably most clearly shows the effects of NP size on caveolae pumping efficiency. Caveolar delivery of NPs displayed an inverse relationship to NP size, with the greatest tissue uptakes occurring for smaller NPs. Caveolae, as 60-80 nm structures, optimally transcytose NP conjugates with HDD less than 35 nm. The 40 nm core Au NPs conjugated to mAPP2 to yield 64 nm HDD complexes still showed high lung uptake ratios relative to nontargeting control but ultimately liver uptake clearly prevailed. To maximize the benefits of caveolae-targeting for NP delivery, our analysis indicates caveolae-based NP delivery systems should focus on NP complexes under 25 nm. As an active transport pathway, targeting caveolae does not require large concentration gradients to move NPs across passive portals and into tissues. Instead, with iv administration of low doses of NPs, caveolae-targeting was able to achieve unprecedentedly high NP levels in lung tissue with minimal accumulation in RES and other off-target-tissues. At least so far, NP composition (i.e., soft vs. hard, dendrimer vs. metal) appears to play a significantly lesser role in targeting compared to size.

RES sequestration of NPs from systemic circulation remains a critical challenge to nano-engineered carriers. NP scavenging by the RES operates faster than passive transvascular delivery to other organs and accounts for NP tropism to organs such as the liver and spleen. Caveolae pumping, in contrast, is robust and fast enough to outcompete RES scavenging by rapidly pulling NPs into target-tissue before they reach RES organs. It likely does so with greater efficiency than NP shielding strategies which only prolong NP circulation time but do not expedite uptake ^5, 48^. Indeed, without conjugation to a CTA, the partially shielded dendrimers in our study were still quickly sequestered by the liver. Caveolae-targeting, in contrast, quickly rerouted smaller NPs to the lungs with little signal in the liver. Not surprisingly, RES scavenging prevails for NP with HDD >60 nm and so near the size and capacity of typical caveolae.

There are other potential factors contributing to nanodelivery to the lungs beyond just the robust pumping power of caveolae. The lungs receive nearly all the cardiac output from the right ventricle and are the first organ contacted after iv injection. This first pass effect and the large microvascular surface area of the lung helps immediate extraction of both retargeted and untargeted NPs. But the small ostia and size of caveolae is likely far more restrictive for NPs than the liver sinusoidal EC gaps (>200 nm)^49^. As NP size increases, rapid entry into caveolae must be diminished which slows immediate blood extraction and provides more opportunity for RES uptake. Future study is needed to discover EC caveolae targets specific for other tissues and then to determine if nano-pumping can still prevail to this same unprecedented degree.

We show here for the first time that NPs when retargeted properly to the CPS can very rapidly and precisely target a single solid tissue; they significantly outperform passive delivery of similarly sized, untargeted controls by as much as two orders of magnitude. The selectivity and enhanced efficiency of the lung CPS was further demonstrated by accumulation of mAPP2-NPs in lung tissue beyond peak blood levels, as shown by a high TCPI. The concentrating power of the caveolae transport system was readily evident within minutes of iv injection and even works against an appreciable concentration gradient, as apparent from the high SUV and TCPI values. CPS-targeting therefore represents a new NP delivery paradigm that can overcome issues of poor tissue penetration and RES sequestration. It promises to enable excellent active and robust tissue-specific penetration and in this case the development of more effective NP-based agents for the imaging, diagnosis, and treatment of many lung diseases including pulmonary fibrosis, acute lung injury, chronic obstructive pulmonary disease, a multitude of lung infections, and pulmonary genetic diseases. Very recently, we showed that an anti-inflammatory therapeutic antibody genetically engineered to target the lung EC CPS inhibited pneumonitis in rats at unprecedentedly low therapeutic doses^32^. By extension, NPs can be loaded with drugs in the future with the now reasonable expectation of using CTAs to enable robust single tissue delivery rendering local therapeutic efficacy while avoiding RES delivery and toxicities.

In this study, we investigated the extent and specificity of caveolae-targeting as a function of NP’s size, a well-known contributor to reduced target tissue uptake and off-target RES accumulation. We characterized trafficking limitations and estimated the optimal diameter range of functionalized NPs for efficient caveolae-targeted delivery. We discovered the unprecedented capability of the CPS to extravasate NPs robustly, quickly and specifically into a single organ, the lung. We quantified for the first time the contrast in how limiting the EC barrier can be to passive entry of NPs into solid, non-RES tissue vs. conversely how enabling ECs can be in tissue-targeted penetration when NPs are properly retargeted via the CPS. How much active extravasation exceeded passive extravasation of NP was quite striking. The 1-2 order in magnitude improvement in lung precision targeting vs similar reduction in RES uptakes were significant. Simply put we have created the first NP to target the CPS and in so doing the first to accumulate more than 50% in a single non-RES tissue. We took NPs normally concentrating in the majority in RES organs within minutes to comprehensively retargeting the majority to the lungs again within minutes But we did discover the CPS has clear size limitations *in vivo* by revealing a remarkable dependence of the lung precision retargeting of NPs on their HDD (12 – 64 nm). NPs with core sizes of 5-20 nm and overall HDDs <35nm targeted the lung within minutes of iv injection and readily achieved robust lung concentrations well beyond peak blood levels (2-4-fold), consistent with caveolae transcytotic pumping against a concentration gradient. RES scavenging of the smaller NPs was rendered minimal by mAPP2 conjugation. All of the control NPs at all sizes tested showed rapid robust RES uptake within minutes. The largest mAPP2 nanocomplexes with HDD >60nm showed lung targeting over controls but mostly accumulated in the liver. Ultimately the greatest precision targeting appears to occur for nanocomplexes below 25nm. These findings will be critical in the design of NP carriers for potential diagnostic imaging and therapeutic applications using active EC transcytosis mediated by the CPS.

## Experimental Methods

### Materials

BioReady™ (10 nm, 20 nm, 40 nm) Carboxyl GNP spheres were sourced from nanoComposix (San Diego, CA). EDC (1-ethyl-3-(3-dimethylaminopropyl)carbodiimide hydrochloride) and Sulfo-NHS (N-hydroxysulfosuccinimide) were purchased from ThermoFisher (USA). Unless otherwise stated all other chemicals were received from Sigma (St Louis, MO).

### Production and purification of antibodies

Recombinant monoclonal antibody APP2 (mAPP2) was genetically engineered in-house as described previously^32^. Non APP2 binding control antibody (mAPP2X), was generated using oligonucleotide site directed mutagenesis to change two amino acids in the middle of heavy chain CDR3. These two changes were sufficient to completely abolish binding to APP2 and hence their lungs targeting potential. Recombinant antibodies transiently expressed in ExpiCHO cells were purified on HiTrap Protein G HP column (Cytiva, Sweden) using AKTA go FPLC platform (GE Healthcare, USA) equipped with an ultraviolet (UV) detector measuring the absorbance at 280 nm.

### Animal use

Animal experiments were carried out in accordance with protocol approved by the Institutional Animal Care and Use Committee (IACUC). Animals were housed in the animal care facility, and those animals which received radiolabeled GNPs were housed and imaged in a separate lead-shielded animal facility. Sprague Dawley rats (females, 100-120 g) were purchased from Envigo (Livermore, CA).

### Biotinylation of PAMAM dendrimer

PAMAM dendrimer G5 (5 mg in 0.05 ml dimethylformamide) was biotinylated with 2 mol eq of NHS-PEG_4_-biotin (ThermoFisher, USA) for 6 h and then primary amines were blocked with 150 mol eq of glycidol for 18 h. Modified dendrimers were then purified by size exclusion chromatography on Sephadex G25. Molecular substitution ratio of biotin was than determined using HABA assay according to manufacturer’s protocol (Pierce, Rockford, IL).

### Radiolabeling of PAMAM dendrimers

PAMAM dendrimer G4 with cystamine core (9.5 μg) or biotinylated G5 with diaminobutane core (13.3 μg) was incubated with 1 mCi of N-succinimidyl-3-(4-hydroxy-3-[^125^I]iodophenyl)propionate in 0.05 ml of dimethylformamide for 1 h at room temperature (RT). Primary amines on the dendrimer were then blocked by glycidol as described above. Afterwards, ^125^I-labeled dendrimer was purified by size exclusion chromatography on Sephadex G25.

### Antibody functionalization of PAMAM dendrons/dendrimers

PAMAM dendrons G4 were produced by reductive cleavage of ^125^I labeled PAMAM dendrimer G4 with cysteamine core using 10 mM Tris(2-carboxyethyl) phosphine (Pierce, Rockford, IL) in PBS for 30 min at RT. PAMAM dendrons were then purified by size exclusion chromatography on Sephadex G25 followed by membrane ultrafiltration (MWCO 10 kDa, Millipore, Billerica, MA). Streptavidin (SAV) (New England BioLabs) was derivatized with sulfosuccinimidyl 4-(N-maleimidomethyl) cyclohexane-1-carboxylate according to manufacturer’s instructions (Pierce, Rockford, IL). PAMAM G4 dendrons with free thiols were then incubated with maleimide-derivatized SAV at a 10:1 molar ratio in the PBS with 5 mM EDTA for 16 h at 4°C. SAV-PAMAM G4 dendron conjugates were purified by ultrafiltration across a molecular weight cut-off membrane of 50 kDa (Millipore, Billerica, MA). SAV-conjugated G4 was linked to biotinylated recombinant APP2-Fab as described previously^30^. Biotinylated ^125^I labeled PAMAM G5 dendrimers were also similarly noncovalently functionalized with trivalent streptabodies via SAV-biotin linkage. See Suppl. Fig. 1.

### GNP-antibody conjugation

GNPs functionalized with PEG12-carboxyic acid (NanoComposix, San Diego, CA) were conjugated with antibodies (mAPP2 or mAPP2X) using EDC and Sulfo-NHS as linking reagents according to a previously described procedure^50^ with appropriate modifications. Briefly, freshly prepared 20 μL of EDC (10 mg/mL) and 40 μL of Sulfo-NHS (10 mg/mL) were first added to 1 mL of BioReady™ GNP solution in microcentrifuge tubes, mixed by vortexing (<20 seconds) and incubated at RT for 30 min under constant rotation. The solution of GNPs with amidated carboxy terminus were subjected to centrifugation (2X) and supernatant having unreacted EDC/sulfo-NHS was carefully removed without disturbing the pellet. The centrifugal parameters of 10 min@3800 CRF, 15 min@10000 CRF and 30 min@20000 RCF were utilized for 40 nm, 20 nm and 10 nm GNPs respectively. The pelleted GNPs resuspended with 1 mL of reaction buffer (5 mM potassium phosphate, 0.5% 20K MW PEG, pH 7.4), were added with 20 μg of antibody (1 mg/mL) and incubated for 10 min at RT under constant rotation. Further incubation of 5 minutes was done after addition of 10 μL quencher (50 % hydroxylamine solution) to deactivate any remaining active NHS-esters. The GNP–antibody suspension was centrifuged (2X) to remove unbound antibody. Final pellet was resuspended with 1 ml of conjugate diluent (0.5X PBS, 1% Tween 20, pH 8) and stored at 4^0^ C.

### Physical, chemical and biological characterization of GNPs

Transmission electron micrographs and core diameter of unconjugated GNPs were acquired at NanoComposix at magnifications ranging from 40 000× to 150 000× using JEOL 1010 transmission electron microscope (JEOL USA, Inc.) operating at 100 KV and were further analyzed by Image J. The size distribution in the form of hydrodynamic diameter (HDD), polydispersity index and zeta potential of both GNPs and antibody conjugated GNPs were measured by dynamic light scattering (DLS) using Zetasizer Nano (Malvern, Worcestershire, UK). Presence of antibody bound to GNPs was confirmed and quantified by BCA protein assay using Pierce™ Micro BCA Protein Assay Kit. The concentration of antibody in conjugated GNPs were adjusted to a range of 7-8 µg/mL prior to their biological characterization. Functionality of antibody bound to GNPs were ascertained by rat APP2 indirect ELISA using the following procedure. 96 well flat-bottom polycarbonate plates (Thermo Scientific, 260836) were coated with 5 ug/ml recombinant rat APP2 (produced in-house) solubilized in 50 mM carbonate buffer at RT overnight. On the following day, plates were washed 3 times using PBS containing 0.05% Tween-20 (PBST). Assay plates were blocked with PBS containing 1% bovine serum albumin and 0.01% Tween-20 for 90 min. Serially diluted antibody or antibody-GNP conjugates were added into the plates and incubated for 2 hr. Afterwards, plates were washed and incubated with anti-hIgG-HRP (1:2000, Jackson, 709-035-149) for 1 hr. After a final series of washes, ABTS substrate (100 μL/well) (KPL, 50-66-01) was added for 15 min and the reaction was terminated using 0.1 M citric acid, pH 4.0 (100 μL/well). The absorbance values were read at 415 nm (Molecular Devices, San Jose, USA) and analysis was done utilizing GraphPad software.

### Radio-iodination of antibodies and conjugation with GNPs

Iodination was carried out using ^125^I radionuclide (Perkin Elmer, IL) and iodination beads according to the manufacture’s protocol (Pierce). Briefly, 2 iodination beads were washed with buffer (5 mM potassium phosphate, pH 7.4), dried with filter paper, and introduced to a reaction vial (Flow Tube) having 300 μL of buffer and Na^125^I (ca. 60 μL, 1 mCi). The mixture was incubated at RT for 5 min with occasional shaking. After 5 minutes, antibody (∼100 μg) in the same buffer was added to the reaction vial, and the mixture was incubated for 15 min at RT. Then, the reaction mixture was passed through PD-10 desalting column (Cytiva) to purify radioimmunoconjugates (mAPP2-^125^I or mAPP2X-^125^I). The concentration of the antibody in radioimmunoconjugates was determined using UV-Visible spectrophotometer (Beckman Coulter). Specific activity (µCi/µg of antibody) of radiolabeled antibodies was calculated from the activity obtained using radio-isotope calibrator (CRC^®^-127R, Capintec) and the molar concentration of the antibody. Radiolabeled antibodies were conjugated with GNPs using the procedure described above.

### Stoichiometry calculations

Molar particle concentrations of GNP radioimmunoconjugates were calculated by taking OD at λ_max_ and comparing with original OD and molar particle concentration of unconjugated GNPs as provided by NanoCompsix. Average number of antibodies per GNP was estimated using both molar particle concentration and specific activity.

### Serum stability study

^125^I-GNP-antibody conjugates were incubated in complete rat serum (ThermoFisher) at 37^0^C for 24 h. Portions of the mixture were sampled at different time points and filtered through Amicon® Ultra-0.5 Centrifugal Filters (MWCO: 100 kDa) purchased from Merck Millipore (Tullagreen, Ireland). The filtrates were collected, and the radioactivity was measured. The percentages of retained (i.e., intact) ^125^I on the GNP conjugates (^125^I-GNP-mAPP2 or ^125^I-GNP-mAPP2X) were calculated using the equation (total radioactivity - radioactivity in filtrate)/total radioactivity.

### Planar and tomographic CT-SPECT imaging

Planar ***γ***-scintigraphic and tomographic CT-SPECT imaging was performed as described in our past work^23^ ^32^ using nanoScan-SPECT/CT preclinical scanner (Mediso, Budapest, Hungary). Briefly, isoflurane-anesthetized healthy SD rats were intravenously (iv) injected with radioimmunoconjugates (∼1.2 µg antibody, ∼10 µCi). Planar and helical CT-SPECT scans were acquired at indicated time points. Region of interest (ROI) analysis of planar images was done using InterView^TM^ FUSION software (Mediso, Budapest, Hungary). SPECT images were acquired using four head gamma cameras equipped M^3^ multi-focus multi-size multi-pinhole whole body high resolution, high sensitivity apertures, APT63 at 60 sec/projection, and the pulse-height analyzer window set at 28.4 keV with a width of 20%. For CT scans, the voltage was set at 50 kV, 300 msec and 720 helical projections. SPECT/CT datasets were reconstructed using Nucline nanoScan software (Mediso, Arlington, VA), further processed and analyzed using VivoQuant software (Invicro, Boston, MA).

### Biodistribution analysis and index calculations

Biodistribution studies and calculations were performed as in our past work^23^ ^26^ ^32^Briefly, at various time points after tail vein injection of radioimmunoconjugates with or without GNPs, the rats were euthanized, and the tissues of interest were dissected, weighed, and radioactivity measured using Gamma Counter (Wallac 1470 Wizard, Perkin Elmer, Finland). Uptake values were calculated as percentage of injected dose per gram of tissue (%ID/g). The immunospecificity index (**ISI**) of targeted radioimmunoconjugates (^125^I-mAPP2 or ^125^I-mAPP2-GNP10/20/40) was estimated by normalizing their tissue uptake (%ID/g) to the tissue uptakes of respective non-targeted radioimmunoconjugates (^125^I-mAPP2X or ^125^I-mAPP2X-GNP10/20/40). The ISI of non-targeted radioimmunoconjugates was estimated by normalizing their tissue uptake with tissue uptake of ^125^I-mAPP2X alone. The tissue targeting index (**TTI**) was calculated as a ratio of tissue to blood uptake in each individual animal and average for the group. Tissue specificity index (**TSI**) was determined as a ratio of TTI of targeted immunoconjugates to the TTI of control non-targeted immunoconjugates. The TSI of non-targeted radioimmunoconjugates was estimated by normalizing their TTI with TTI of ^125^I-mAPP2X alone. Specific uptake value (**SUV**) was calculated as tissue uptake normalized to the uptake that would be reached after homogeneous penetration throughout all tissues. Tissue concentrating power index (**TCPI**) was estimated as a ratio of uptake in each tissue to the peak blood level. N=3 for all experimental groups. All results were expressed as mean ± standard deviation (S.D.).

### Inductively coupled plasma mass spectrometry (ICP-MS)

Healthy SD rats in group (n=3) were injected iv with immunoconjugates of mAPP2 and mAPP2X along with their GNP10 and GNP40 derivatives. Animals were euthanized and tissues of interest were harvested 1 h post injection time point. Known mass of each sample were taken up in Savillex vials having 5 ml of freshly prepared aqua regia (1:3 ratios of concentrated nitric acid and hydrochloric acid) and heated overnight at 60 °C for complete digestion. After samples were fully dissolved, 40 μl of the digest were diluted with deionized water to 10 mL in 15 ml centrifuge tubes and analyzed at the Environmental and Complex Analysis Laboratory (ECAL) of University of California, San Diego on iCAP™ RQ ICP-MS (ThermoFisher) instrument and referenced to a HAuCl4-derived calibration.

### Statistical methods

Data are expressed as mean +/-standard deviation of replicate experiments. The number of measurements is given in each figure legend. Group sizes were determined with the goal of ensuring a minimal power of 0.80 to detect expected effect sizes of .5 to 1.0 between controls and other groups at a two-sided a of 0.05. We have utilized general linear models (GLMs) followed by post-hoc multiple comparison procedures if warranted as our standard method of analysis, on untransformed data. We assessed normality on the residuals from the GLM fits with Lilliefors-corrected Kolmogorov-Smirnov tests; invariably, p-values were nonsignificant at the a=0.05 level. Analyses were performed in R v4.2.1 (R Core Team, 2021) and Prism 10 (Graphpad, 2023). Significance was set conventionally at p<0.05.

## Abbreviations

APP2: aminopeptidase P2
CTA: caveolae-targeting antibody
SPECT-CT: computed tomography co-registered with single-photon emission computerized tomography
%ID/g: percent of injected dose per gram of tissue
EC: endothelial cell
mAb: monoclonal antibody
rAPP: recombinant APP
RES: reticulo-endothelial system
NP: nanoparticle
GNP: gold nanoparticles
mAPP2: APP2-specific chimeric human/mouse monoclonal antibody
RINC: radioimmunonanoconjugate
HDD: hydrodynamic diameter
ROI: region of interest
PAMAM: poly(amidoamine)
CPS: caveolae pumping system

---

**Supplement Fig. 1.**
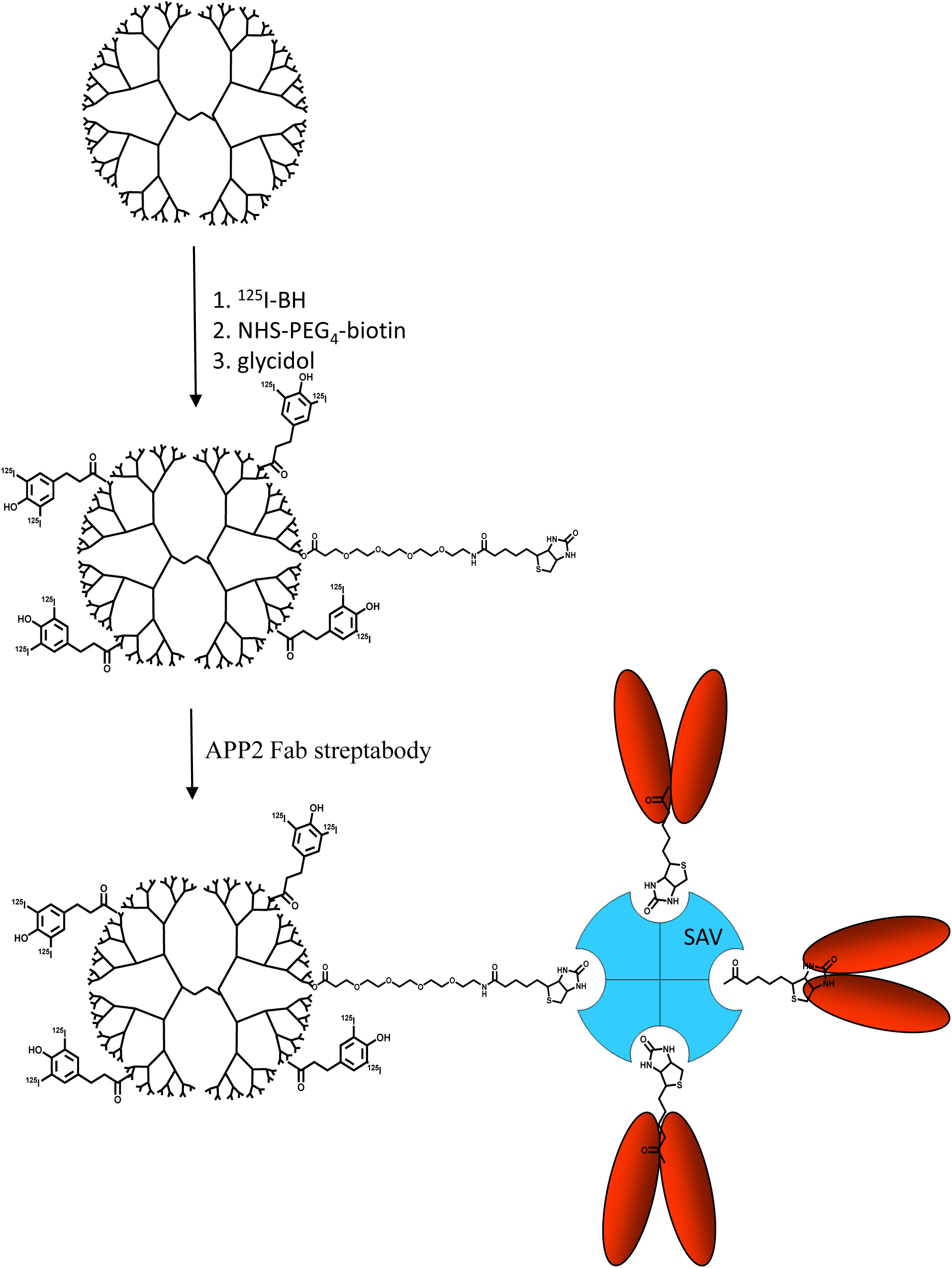
Functionalization of PAMAM G5 dendrimers with APP2 Fab to form supramolecular nanoassembly. PAMAM G5 were biotinylated using NHS-PEG4-biotin then radiolabeled with N-succinimidyl-3-(4-hydroxy-3-[_125_I]iodophenyl)propionate and terminated with glycidol. To form supramolecular nanoassembly, the biotinylated _125_I labeled PAMAM G5 dendrimers were then noncovalently functionalized with trivalent APP2 streptabodies via streptavidin (SAV)-biotin linkage.

**Supplement Fig. 2.**
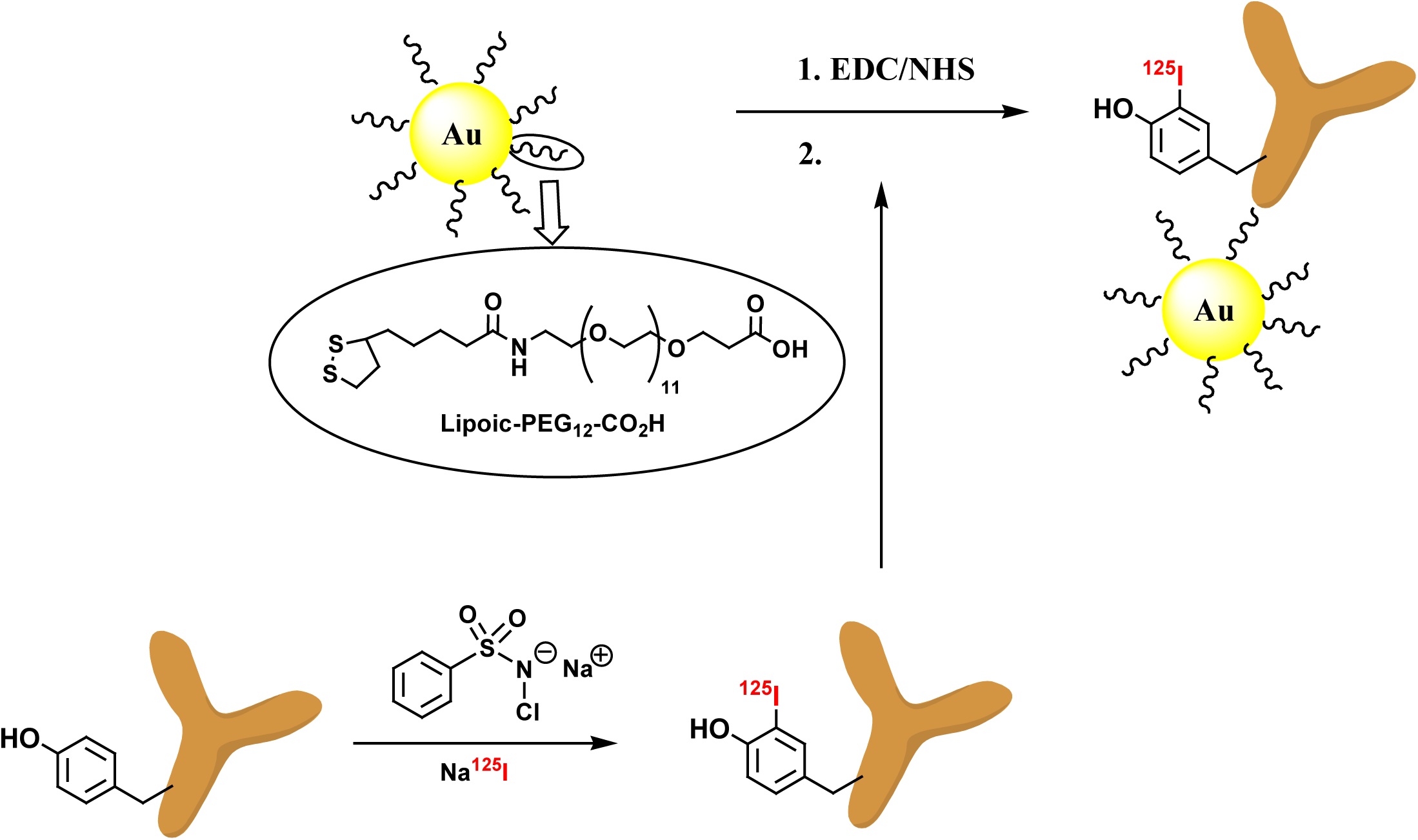
Functionalization of gold nanoparticles with radioiodinated mAPP2. PEG-Carboxilic acid terminated gold nanopartilces were conjugated to mAPP2 antibody as described in Methods. EDC: 1-ethyl-3-(3-dimethylamino) propyl carbodiimide, NHS: N-hydroxysulfosuccinimide.

**Supplement Fig 3.**
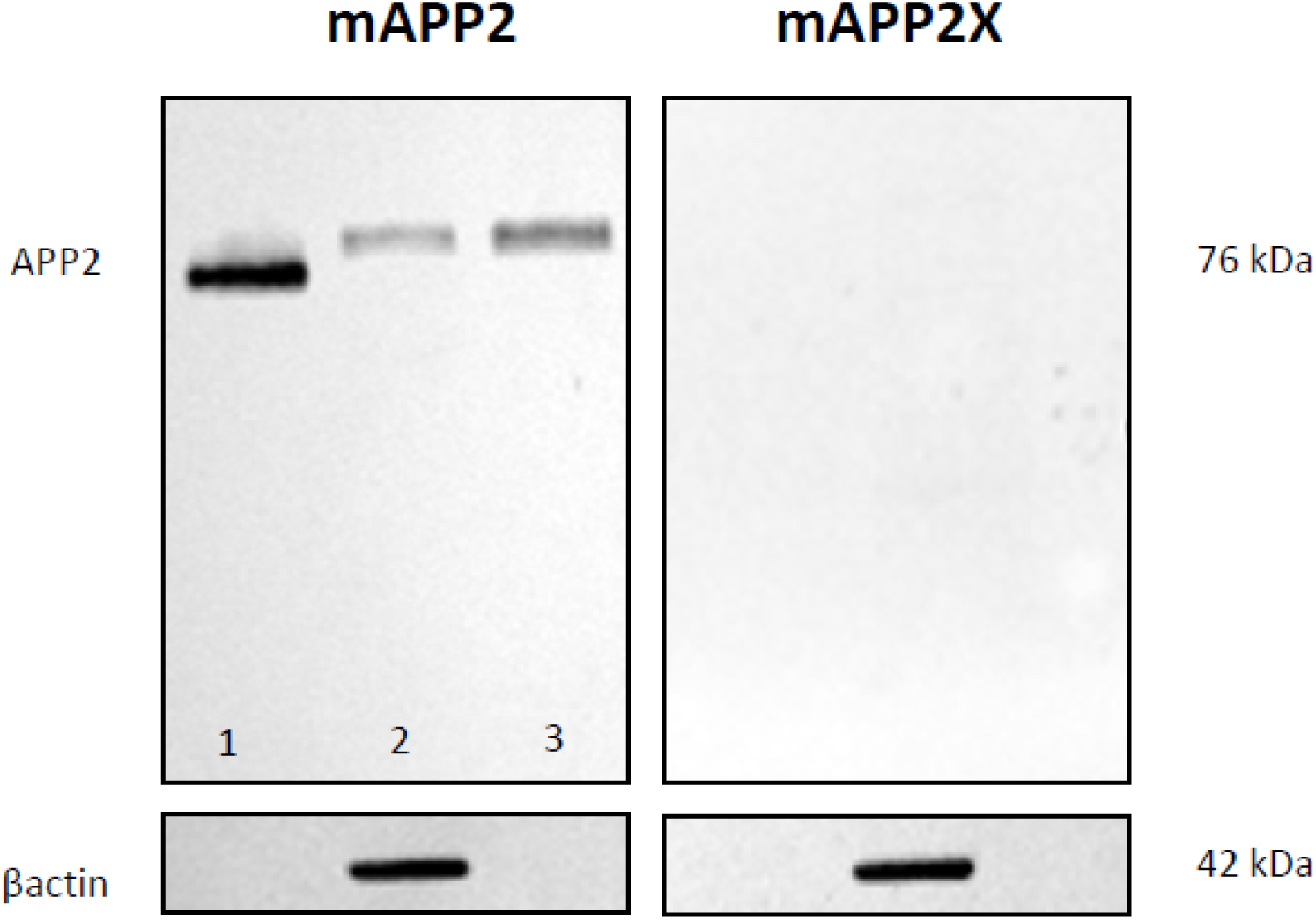
Western blot validation of antibody specificity towards rat APP2. Membranes, containing (1) 50 ng recombinant rat APP2 protein (aa2-648 + 6XHIS), (2) 30 µg rat lung homogenate, and (3) 30 µg purified plasma membrane fraction of rat lung-derived endothelial cells, probed with mAPP2 and non-specific, mutated mAPP2X show distinct difference in target recognition. mAPP2X shows no binding to rat APP2 nor any native rat lung proteins.

